# Primary Cilium-dependent Humoral Bioactive Factors Acts in a Paracrine Manner to Control Fibroblast Cell Migration

**DOI:** 10.1101/2025.08.20.671189

**Authors:** Faryal Ijaz, Koshi Imami, Koji Ikegami

**Author notes:** Co-corresponding author: Faryal Ijaz, Department of Anatomy and Developmental Biology, Graduate School of Biomedical and Health Sciences, Hiroshima University, 1-2-3 Kasumi, Minami-Ku, Hiroshima 734-8553, Japan; Koji Ikegami, Department of Anatomy and Developmental Biology, Graduate School of Biomedical and Health Sciences, Hiroshima University, 1-2-3 Kasumi, Minami-Ku, Hiroshima 734-8553, Japan.

## Abstract

The primary cilium is classically recognized as a signal-reception hub, yet its role in mediating cell-to-cell communication via signal spread remains poorly defined. Here, we uncover a previously unrecognized paracrine function of the primary cilium in fibroblasts. Conditioned medium from ciliated wild-type NIH/3T3 cells enhanced wound healing in primary cilium-deficient fibroblasts, in contrast to conditioned medium derived from primary cilium-deficient NIH/3T3-*Kif3a*-KO or NIH/3T3-*Dync2h1*-KO cells. Fractionation of conditioned medium revealed that the wound healing activity resided predominantly in the 100K × g soluble supernatant (Sup-100K), rather than in extracellular vesicle (EV) fractions. Untargeted metabolomic analysis identified lysophosphatidylcholine (LPC) (14:0) as a key bioactive metabolite enriched in WT-Sup-100K secretome. Supplementation of LPC(14:0) restored wound healing capacity in NIH/3T3-*Kif3a*-KO cells to levels comparable to WT-Sup-100K treatment. Transcriptomic profiling of target cells revealed that WT-Sup-100K upregulated expression of extracellular matrix (ECM)-associated genes, including *Ogn*, *Igf2*, and *Mfap4*, while EVs modestly enhanced early ECM remodeling via induction of *Nid2*. Together, these findings demonstrate that the primary cilium coordinates a wound healing secretome in fibroblasts through the regulated release of LPC(14:0) and other soluble factors that activate ECM-remodeling pathways in recipient cells. This work expands the functional repertoire of the primary cilium and establishes its critical role in coordinating paracrine regenerative signaling.

## Introduction

A primary cilium is a specialized organelle that extends from the surface of most mammalian cells and plays an essential role in the detection of extracellular signals. This solitary antenna-like structure is fundamental for cellular signal transduction, facilitating a wide range of developmental and physiological processes. These processes include sensory functions, signal transduction, cell cycle regulation, embryonic development, tissue homeostasis, cell migration and differentiation, planar cell polarity, mechanosensation, chemosensation, photoreception, and olfaction. The presence and proper functioning of primary cilia are essential for normal development and physiology across diverse tissues and organs in the body. Defects in primary cilia structure or function can lead to a wide range of developmental abnormalities and diseases collectively known as ciliopathies^[1]^. These disorders can affect multiple organ systems and manifest with diverse phenotypes such as cystic kidneys, retinal degeneration, skeletal malformations, brain abnormalities, obesity, and polydactyly.

The primary cilium hosts signaling components such as Hedgehog, Wnt, and platelet-derived growth factor receptor alpha (PDGFRα), thereby functioning as a central hub for interpreting both chemical and mechanical signals from the external environment^[2–5]^. When receptors in the cilium are activated, they employ secondary messengers such as calcium and cAMP to either directly influence downstream signaling mediators or initiate signaling via posttranslational modifications (PTMs)^[6]^. This mechanism is predominantly associated with GPCRs^[7–12]^. Conversely, other types of ciliary chemoreceptors, including Receptor Tyrosine Kinases (RTKs) and Transforming Growth Factor beta (TGFβ) receptors, directly phosphorylate their respective targets, altering ciliary signaling dynamics and prompting downstream cellular responses^[13]^. Additionally, the specific lipid compositions of ciliary membranes, along with associated protein complexes, significantly contribute to complex signaling pathways that facilitate the transmission of diverse signals to the cells^[10]^.

Beyond their role in sensing incoming mechanical and chemical signals, primary cilia function as a means of releasing bioactive factors into the environment. The ability of cilia to emit ectosomes into the extracellular environment has been observed in diverse organisms. This phenomenon has been documented in Chlamydomonas^[14–18]^, C. elegans^[19]^and mammalian species^[20,21]^. In mammalian cells, the release of ectosomes from primary cilia appears to be essentially the result of the cutting of the tips of the cilia^[20,21]^. The extracellular vesicles (EVs) released from primary cilia contain proteins and mRNAs and are thought to be bioactive^[21–25]^. Several studies have demonstrated the bioactive properties of ciliary EVs, which play a crucial role in modulating cellular signaling pathways during development and disease conditions^[26,27]^. The proliferation of glioma cells increased when they were co-incubated with culture medium containing primary ciliary EVs from cancer cells. This effect was confirmed by comparing it with culture media from cells that lacked primary cilia^[28]^. In bbs4−/− and bbs8−/− mutants, where intraflagellar transport is disrupted, certain proteins that were initially intended for release via large ciliary EVs were instead found to be enriched in small EVs originating from the cell cytoplasm^[26]^. These EVs, which contained both proteins and miRNAs, were capable of influencing Wnt signaling in recipient cells^[26]^. Furthermore, primary cilia can relay signals through secretion of bioactive soluble factors^[29,30]^. Osteocytes suppress cancer cell proliferation and increase migration via tumor necrosis factor alpha (TNF-α) secretion and this action is regulated by osteocyte primary cilia^[30]^.

Lipids are increasingly recognized as critical biological factors in cellular signaling that influence a wide array of physiological and pathological processes^[31,32]^. Certain bioactive lipids, including lysophosphatidylcholine (LPC), lysophosphatidic acid (LPA), and sphingosine-1-phosphate (S1P), act as signaling molecules that regulate essential cellular functions such as cell migration, proliferation, and apoptosis^[33–35]^. Among these, LPCs have garnered significant attention owing to their diverse biological roles and involvement in disease development. LPC is produced through the enzymatic cleavage of phosphatidylcholine by phospholipase A2^[36]^ and is a major component of oxidized low-density lipoprotein, which is a key mediator of atherosclerosis^[37]^. Beyond its role in cardiovascular diseases, LPC has also been implicated in modulating immune responses, oxidative stress, and cancer cell metastasis^[38–41]^. LPC exerts its effects through various mechanisms, including activation of G-protein-coupled receptors, modulation of ion channels, and interactions with membrane lipids^[42–45]^.

Most investigations of primary cilia have focused on their roles in sensing extracellular signals and activating intracellular pathways that regulate cellular functions, particularly in mammalian cells. While these studies have deepened our understanding of how primary cilia function within individual cells, less attention has been given to how primary cilia influence other cells through secreted or surface-presented factors. This study aimed to address this gap by examining how primary cilia affect the release or display of extracellular signals that modulate the behavior and function of surrounding or distant cells. By exploring these primary cilium-mediated intercellular interactions, we sought to uncover new mechanisms by which ciliary signaling contributes to communication between cells. We found that primary cilia regulate the secretion of bioactive extracellular lipid molecules that, in turn, control fibroblast migration and proliferation through humoral cell-to-cell signaling.

## Materials and Methods

### Cell Culture

All the cell lines were maintained at 37°C with 5 % CO_2_. NIH/3T3 cells were obtained from ATCC (CRL-1658) and cultured in Dulbecco’s modified Eagle’s medium (DMEM)-high glucose (044-29765, Wako) supplemented with 10 % fetal bovine serum (FBS; Gibco). The NIH/3T3-*Kif3a*-KO and NIH/3T3-*Dync2h1*-KO cell lines were established as previously described^[21,46]^. For cilia formation, the medium of the confluent cells was replaced with serum-reduced DMEM-high glucose supplemented with 1 % FBS for 12 h. To release extracellular humoral factors into the medium, cells were first cultured in DMEM-high glucose supplemented with 10 % FBS for 72 h and then in serum-reduced DMEM-high glucose supplemented with 0.2 % FBS for 15 h.

### Reagents

The primary antibodies used in this study were as follows: ARL13B (mouse mAb N295B/66; ab136648; Abcam), ARL13B (rabbit pAb; 17711-1-AP; Protein Tech), and anti-γ-tubulin (mouse mAb GTU-88; T6557; Sigma-Aldrich, St Louis, MO, USA). Alexa fluorophore-conjugated secondary antibodies (Thermo Fisher Scientific) were used for immunofluorescence microscopy. The 14:0 form of LPC (Avanti Polar Lipids, USA) was dissolved in methanol (50 mM stock solution). Click-iT™ EdU Alexa Fluor™ 555 Cell Proliferation Kit for Imaging (Invitrogen, C10338).

### Immunocytochemistry (ICC)

Cultured cells were fixed with 4 % paraformaldehyde (PFA, pH 7.5) for 30 min at 37°C. The cells were washed with PBS, blocked, and permeabilized with 5 % normal goat serum containing 0.1 % Triton X-100 in PBS for 1 h at room temperature, followed by incubation with primary antibodies for 24 h at 4°C. After washing in PBS, the cells were incubated for 2 h at room temperature with Alexa Fluor-conjugated secondary antibodies and DAPI (1:1000; DOJINIDO). Images were acquired using a confocal microscope (Olympus FV1000) equipped with an oil-immersion lens (60×, NA 1.35).

### Conditioned Medium, Supernatant and EVs Fractions Preparation

The FBS used for humoral bioactive factor collection was pre-depleted via overnight centrifugation at 100,000 × g to remove tiny FBS-intrinsic vesicles, exosomes, and ectosomes. Wild-type, *Kif3a*-KO or *DynC2h1*-KO NIH/3T3 cells were grown to confluence. The culture medium was replaced with a fresh medium containing 0.2 % FBS. The conditioned culture media were collected 15 h later. Humoral bioactive factors released from cultured cells into the media were collected using a three-step centrifugation process. The culture media were first centrifuged at 2,000 × g for 20 min to remove large cell debris. The supernatant from the first centrifuge was further centrifuged at 10,000 × g for 30 min. At this step, conditioned media (CMs) were either used as it was or ultracentrifuged for further fractionation. For further fractionation, the supernatant of the second centrifuge was centrifuged at 100,000 × g for 3 h using an MLA-55 angle rotor (Beckman). Supernatant fractions depleted of EVs were collected following ultracentrifugation. The EV pellets left after the separation of supernatant fractions were resuspended in PBS to remove residual aggregates of macromolecules and ultracentrifuged once again at 100,000 × g for 1 h. After the second ultracentrifugation step, EV pellets were resuspended in convenient volumes of DMEM-high glucose supplemented with 0.2 % pre-depleted FBS. In parallel, the supernatant of the second centrifuge was concentrated 40–50 times by centrifugation at 4000 × g at 4°C using Amicon Ultra 15 ml filters (Millipore, Burlington, MA, USA) with a 100 kDa molecular weight cut-off (MWCO). The volume was made up to 1 mL using DMEM-high glucose. The CM, supernatant, and EV pellet were then subjected to a wound healing assay, proliferation assay and omics.

### Wound Healing Assay

CMs, supernatant, and EVs collected from wild-type, *Kif3a*-KO, and *Dync2h1*-KO NIH/3T3 cells were used to compare the effects of these fractions on the wound healing capacity of target cells. NIH/3T3 *Kif3a*-KO cell line was seeded with 4 × 10^5^ cells/well to confluency to a 12-well plate (Thermo Fisher Scientific) in a 1 mL culture medium and incubated for 24 h. After 24 h, a gap was created by straight scratching using 200 μL pipette tips. The cell medium was then removed to clear dead and de-attached cells, and the culture media were replaced with 1 mL of CMs, supernatant, or EVs fractions. The treatment medium was replaced every 12 h for 48 h. Finally, 0th, 12th, 24th, 36th and 48th-hour images of the cells were taken. ImageJ’s “MRI Wound Healing Tool” plugin was employed to analyze and quantify the closure rates of the created wounds, and the averaged distance recovered was measured in pixels.

### Proliferation Assay

Supernatant collected from wild-type and *Kif3a*-KO NIH/3T3 cells were used to compare the effects of these fractions on the proliferation capacity of target cells. NIH/3T3 *Kif3a*-KO cell line was seeded with 4 × 10^5^ cells/well to confluency to a 12-well plate (Thermo Fisher Scientific) in a 1 mL culture medium and incubated for 24 h. After 24 h, a gap was created by straight scratching using 200 μL pipette tips. The cell medium was then removed to clear dead and de-attached cells, and the culture media were replaced with 1 mL of supernatant fractions containing EdU. The treatment medium was replaced every 12 h for 48 h and then fixed and permeabilized. EdU was used to visualize proliferating cells using Click-iT™ EdU Alexa Fluor™ 555 Imaging kit. Cells were counter stained with 1 μg/mL DAPI (DOJINDO). Finally, cells were imaged on LEICA DMI 3000B equipped with K8 camera (LEICA). ImageJ was used to count the EdU positive and total number of cells in the wound and non-wound area.

### Proteomic Profiling

EVs collected from wildtype, *Kif3a*-KO, NIH/3T3cell lines were used to compare the effects on gene expression profile of recipient cells. NIH/3T3 *Kif3a*-KO cell line was seeded with 4 × 10^5^ cells/well to confluency to a 12-well plate (Thermo Fisher Scientific) in a 1 mL culture medium and incubated for 24 h. After 24 h, the culture media were replaced with 1 mL of the EVs fractions. After 12 h, treated cells were collected for proteomic analysis. Cell pellets were resuspended with 100 mM Tris-HCl pH 9.0 containing 8 M urea. Proteins were reduced with 10 mM dithiothreitol (DTT) (FUJIFILM Wako) for 30 min at 37°C, followed by alkylation with 50 mM 2-iodoacetamide (IAA) (FUJIFILM Wako) for 30 min at room temperature in the dark. The proteins were digested with 1 μg lysyl endopeptidase (LysC) (FUJIFILM Wako) for 3 h at 37°C. The samples were diluted to 2 M urea with 50 mM ammonium bicarbonate. The proteins were further digested with 1 μg trypsin (Promega) overnight at 37°C on a shaking incubator. The resulting peptides were acidified with 0.5% TFA (final concentration), and purified a StageTip containing SDB-XC (upper) and SCX (bottom) Empore disk membranes (GL Sciences) as described before ^[47,48]^

Nano-scale reversed-phase liquid chromatography coupled with tandem mass spectrometry (nanoLC/MS/MS) was performed by an Orbitrap Fusion Lumos mass spectrometer (Thermo Fisher Scientific) connected to a Thermo Ultimate 3000 RSLCnano pump and an HTC-PAL autosampler (CTC Analytics, Zwingen, Switzerland) equipped with a self-pulled analytical column (150 mm length × 100 μm i.d.) packed with ReproSil-Pur C18-AQ materials (3 μm, Dr. Maisch GmbH, Ammerbuch, Germany). The mobile phases consisted of (A) 0.5% acetic acid and (B) 0.5% acetic acid and 80% acetonitrile (ACN). Peptides were eluted from the analytical column at a flow rate of 500 nL/min with the following gradient: 5–10% B in 5 min, 10–40% B in 60 min, 40–99% B in 5 min, and 99% for 5 min. The Orbitrap Fusion Lumos instrument was operated in the data-dependent mode with a full scan in the Orbitrap followed by MS/MS scans for 3 s using higher-energy collisional dissociation (HCD). The applied voltage for ionization was 2.4 kV. The full scans were performed with a resolution of 120,000, a target value of 4 × 10^5^ ions, and a maximum injection time of 50 ms. The MS scan range was m/z 300–1,500. The MS/MS scans were performed with a 15,000 resolution, a 5 × 10^4^ target value, and a 50 ms maximum injection time. The isolation window was set to 1.6, and the normalized HCD collision energy was 30. Dynamic exclusion was applied for 20 s.

All raw data files were analyzed and processed by MaxQuant (v1.6.5.0),^[49]^and the database search was performed against the mouse UniProt database (version 2019-04) spiked with common contaminants and enzyme sequences. Search parameters included two missed cleavage sites and variable modifications such as methionine oxidation, protein N-terminal acetylation and deamidation of glutamine and asparagine. Cysteine carbamidomethylation was set as a fixed modification. The peptide mass tolerance was 4.5 ppm, and the MS/MS tolerance was 20 ppm. The false discovery rate (FDR) was set to 1% at the peptide spectrum match (PSM) level and protein level. For protein-level quantification in all experiments, ‘unique + razor’ peptides were used. For label-free quantification, a minimum of two ratio counts was used for quantification. Only proteins quantified in 2 out of the 3 replicates in at least one condition were used for further analysis, and missing values were imputed from a normal distribution of log2 LFQ intensity using a default setting (width 0.3, down shift 1.8) in Perseus (v.1.6.5.0)^[50]^.

### RNA-Seq and Analyses

*Kif3a*-KO cells were seeded with 4×10^5^ cells/well to confluency to a 12-well plate (Thermo Fisher Scientific) in a 1 mL culture medium and incubated for 24 h. After 24 h, culture media were replaced with 1 mL of supernatant fractions derived from wild-type and *Kif3a*-KO NIH/3T3 cell lines and replaced every 12 h. After 24 h and 48 h of treatment, cells were collected, and total RNA was extracted using the RNeasy Mini Kit (Qiagen, Germany) and sequenced. Sequenced readings for each sample were aligned to mm39 using HISAT and Bowtie2, and differentially expressed genes (DEGs) were identified using BGI’s Dr. Tom System. KEGG enrichment analyses were done using ShinyGO 0.82^[51]^ (https://bioinformatics.sdstate.edu/go/).

### Lipidomics of Cell Secretome and Validation by Rescue Experiments

Supernatant fractions obtained from ultracentrifugation of CMs derived from the wild-type, *Kif3a*-KO, and *Dync2h1*-KO NIH/3T3 cell lines were subjected to lipidomic analysis. A Waters UPLC I-Class Plus (Waters, USA) tandem Q Exactive high-resolution mass spectrometer (Thermo Fisher Scientific, USA) was used for separation and detection of metabolites. Chromatographic separation was performed on a CSH C18 column (1.7 µm 2.1*100 mm, Waters, USA). Data acquisition and processing were performed using LipidSearch v.4.1 (Thermo Fisher Scientific, USA) and MetaX, respectively.

Rescue experiments were performed to validate the role of the identified lipids in cell migration. Briefly, a wound healing assay was performed in which recipient cells were treated with supernatant fractions derived from wild-type and *Kif3a*-KO cell lines as controls and supernatant fraction obtained from *Kif3a*-KO cell lines supplemented with 10 µM LPC(14:0) as treatment.

### Statistical Analyses

Statistical analyses and graphing were performed using GraphPad Prism 10.0 software (GraphPad Software, Inc.) and Excel. Pairwise comparisons of wound healing were performed using the t-test. Data are presented as mean±standard error of the mean (SEM) of three or four independent analyses. Statistical significance was set at a *p*-value < 0.05.

## Results

### Conditioned medium derived from ciliated wild-type NIH/3T3 cells promote wound healing more than that from primary cilium-deficient knockout cells

To explore whether cilia-dependent humoral effects are observed, we first investigated the effect of ciliated fibroblast-conditioned media (CM) on cilia-depleted fibroblast cell behavior. To this end, we compared NIH/3T3-WT CM with CM from cultures of primary cilium-deficient cell lines lacking KIF3A (NIH/3T3-*Kif3a*-KO), a protein dispensable for ciliogenesis. The absence of primary cilia was confirmed by immunostaining of a primary cilium marker, ARL13B (Figure 1A). The CM used was collected through a two-step centrifugation process with dead cells and debris removed, and the resulting supernatant after 10K × g centrifugation was referred to as conditioned medium 10 K (CM-10K) (Figure 1B). Primary cilia-deficient NIH/3T3-*Kif3a*-KO cells were exposed to either of these CM-10K media after wounding. The wound healing rate of the target NIH/3T3-*Kif3a*-KO cells was significantly enhanced in the CM-10K obtained from ciliated WT cell cultures compared to CM-10K collected from primary cilia-deficient *Kif3a*-KO cell cultures (Figure 1C; dark grey boundary). Cells exposed to WT conditioned medium had an average wound recovery rate of approximately 56 % at 48 h, whereas cells exposed to *Kif3a*-KO conditioned medium had a wound recovery rate of 39 % at 48 h (Figure 1D). The target cells showed minimal recovery in a fresh culture medium; the wound recovery rate was ∼10 % (Figure 1C-D). We also measured the actual recovery level as averaged distance recovered to quantify and compare more precisely the level of wound healing (Figure 1E). The averaged distance recovered of the target NIH/3T3-*Kif3a*-KO cells after 48 h of wounding was almost twice in the WT conditioned medium compared to that of target cells exposed to conditioned medium from *Kif3a*-KO cells (Figure 1E; ∼12 px versus ∼7 px with *p* = 0.0408).

**Figure 1:**
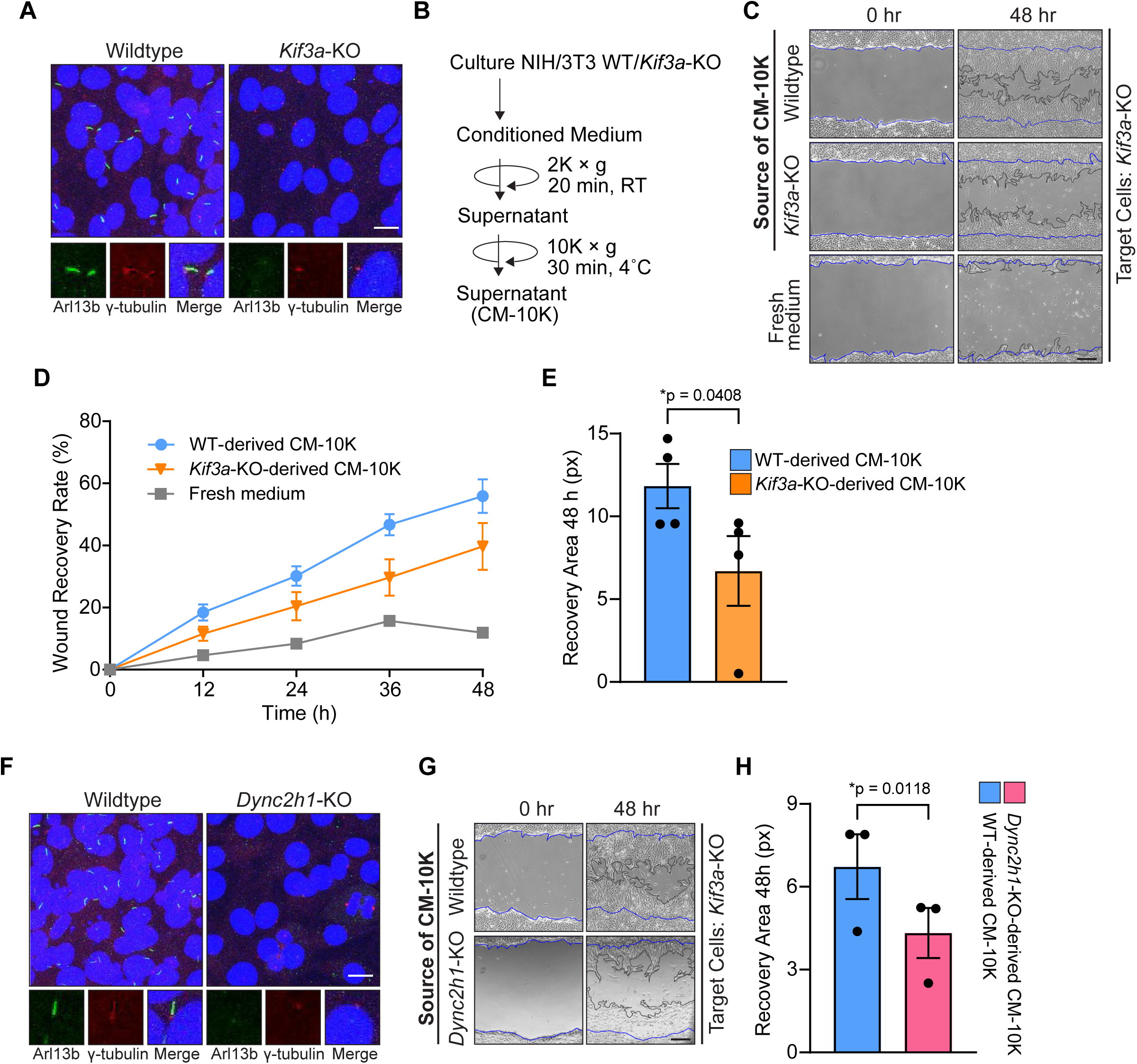
Wild-Type NIH/3T3 conditioned medium facilitated primary cilium-dependent wound healing. (**A**) Fluorescence microscopy images after immunocytochemistry (ICC) showing cells with primary cilium stained for ciliary marker (ARL13B; green), centriole marker (γ-tubulin; red), and nuclei (DAPI; blue) in NIH/3T3-WT and NIH/3T3-*Kif3a*-KO cells. Scale bar, 10 μm. (B) Schematic representation of fractionation of the supernatant (CM-10K) from the conditioned medium. (C) Microscopic images of wound closure in untreated (0.2 % FBS, bottom panel) and treated with WT-CM-10K or *Kif3a*-KO-CM-10K (upper and middle panels) *Kif3a-KO* cells at 0 and 48 h after wound creation. Blue lines define the area covered by the cells at 0 h after treatment. Dark-grey lines define the area covered by the cells after 48 h. Scale bars, 250 μm. (D) Quantification of the wound area recovery rate of *Kif3a-KO* cells at 12h, 24h, 36h and 48 h untreated (light grey) and treated with WT-CM-10K or *Kif3a*-KO-CM-10K (blue and orange) presented as percentages (%). (E) Quantitative analysis of the wound healing assay as the recovery area in pixels (px) at 48 h from C. (F) Fluorescence microscopy images after immunocytochemistry (ICC) showing cells with primary cilia stained for ciliary marker (ARL13B; green), centriole marker (γ-tubulin; red), and nuclei (DAPI; blue) in NIH/3T3-WT and NIH/3T3-*Dync2h1*-KO cells. Scale bar, 10 μm. (G) Microscopic images of wound closure of *Kif3a-KO* cells treated with WT-CM-10K or *Dync2h1*-KO-CM-10K (upper and middle panels) at 0 and 48 h after wound creation. Blue lines define the area covered by the cells at 0 h after treatment. Dark-grey lines define the area covered by the cells after 48 h. Scale bars, 250 μm. (H) Quantitative analysis of the wound healing assay from G. All data are presented as the mean ± SEM (n = 3–4 per group); \**p* < 0.05.

We further verified that WT conditioned medium promoted wound healing by using another primary cilium-depleted cell line that lacked DYNC2H1 (NIH/3T3-*Dync2h1*-KO), a protein involved in IFT transport. The deficiency of primary cilia was also confirmed by immunostaining of a primary cilium marker, ARL13B (Figure 1F). The recovery area of the target NIH/3T3-*Kif3a*-KO cells after 48 h of wounding was larger in the WT-CM-10K (Figure 1G; upper panel; dark grey boundary) compared to the recovery area of the target cells exposed to CM from *Dync2h1*-KO cells (Figure 1G; bottom panel; dark grey boundary). In the quantification, the averaged distance recovered of the target NIH/3T3-*Kif3a*-KO cells after 48 h of wounding was ∼1.7-fold in the WT conditioned medium compared to that of target cells exposed to conditioned medium from *Dync2h1*-KO cells (Figure 1H; ∼7 px versus ∼4 px with *p* = 0.0118). These results further support a scenario that NIH/3T3-WT CM promotes wound healing of cilia-deficient *Kif3a*-KO cells in a primary cilium-dependent manner.

### 100K × g supernatant of conditioned medium derived from ciliated wild-type NIH/3T3 cells promotes migration but not proliferation in primary cilium-deficient target cells

The result that NIH/3T3-WT CM shows better wound healing of cilia-depleted target cells than CMs from primary cilium-deficient cells suggests that the NIH/3T3-WT CM contains humoral bioactive factors that exert the promotion of wound healing. Humoral bioactive factors are released from cells either in the form of EVs or as soluble biomolecules in the form of proteins, nucleic acids, or lipids. We thus investigated which EVs or soluble fraction of the NIH/3T3-WT CM promoted wound healing of target *Kif3a*-KO cells. To separate the EV pellet fraction (EV-P3) and the 100K × g supernatant fraction (Sup-100K), CM-10K prepared from NIH/3T3-WT CM was further ultracentrifuged (Figure 2A). Effects of these two fractions on wound healing in target *Kif3a*-KO cells were investigated with EV-P3 resuspended in fresh culture medium. The recovery from wound of *Kif3a*-KO cells exposed to Sup-100K prepared from NIH/3T3-WT CM (WT-Sup-100K) appeared to be comparable to that exposed to WT CM-10K (Figure 2B; top and middle panel; dark grey boundary) and better than that exposed to resuspended EV-P3 (Figure 2B; bottom panel; dark grey boundary). Quantitative analysis of wound recovery rates over time corroborated these findings. At 12, 24, 36, and 48 hours post-treatment, the recovery area in WT-Sup-100K-treated cells closely mirrored the WT-CM-10K group (13 % vs. 18 %, 27 % vs. 30 %, 44 % vs. 46 %, and 47 % vs. 55 %) (Figure 2C). In contrast, the WT-EV-P3 fraction elicited progressively lower recovery, reaching only 41% at 48 h. In the early phase of healing (12 and 24 h), the wound closure rates were comparable between WT-Sup-100K and WT-EV-P3 treatments (13 % vs. 13 % and 27 % vs. 25 %) suggesting an initial contribution of EVs to migratory activity. However, this effect waned by 48 h, at which point the recovery rate in WT-EV-P3-treated cells became indistinguishable from that of cells treated with CM-10K derived from cilium-deficient *Kif3a*-KO cells (Figure 2C), indicating that soluble factors in WT-Sup-100K are primarily responsible for sustained wound healing.

**Figure 2:**
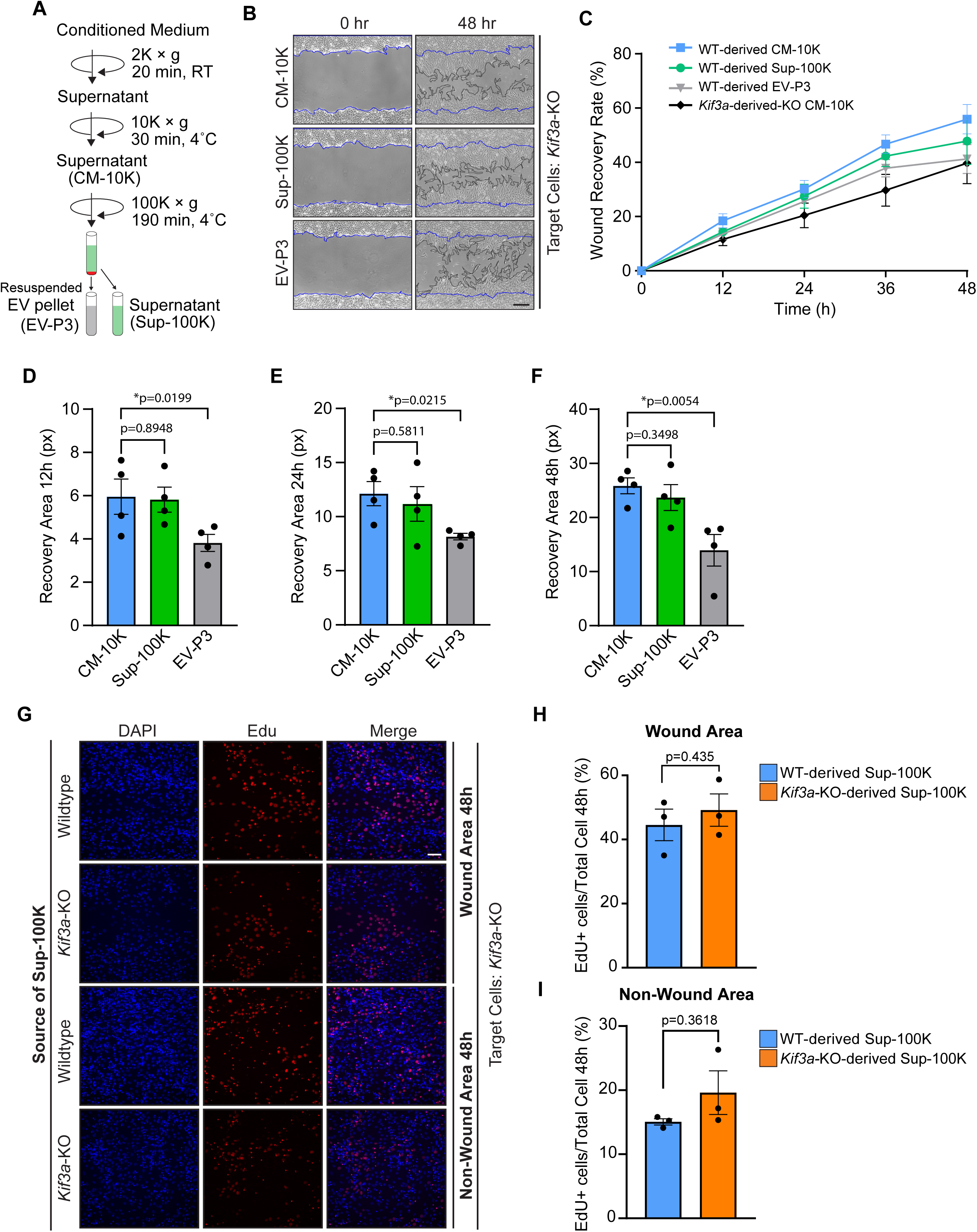
The 100K × g supernatant fraction derived from wild-type NIH/3T3 conditioned medium facilitates wound healing in target cells. (**A**) Schematic representation of fractionation of the supernatant (Sup-100K) from conditioned medium after ultracentrifugation. (B) Microscopic images of wound closure in *Kif3a*-KO cells treated with WT-CM-10K (upper panel), WT-Sup-100K (middle panel), and WT-EV-P3 (lower panel) at 0 and 48 h after wound creation. Blue lines define the area covered by the cells at 0 h after treatment. Dark-grey lines define the area covered by the cells after 48 h. Scale bars, 250 μm. (C) Quantification of the wound area recovery rate of *Kif3a-KO* cells at 12h, 24h, 36h and 48 h after treatment with WT-CM-10K (blue), *Kif3a*-KO-CM-10K (black), WT-Sup-100K (green), and WT-EV-P3 (grey) presented as percentages (%). (D) Quantitative analysis of the wound-healing assay as the recovery area in pixels (px) at 12 h. (E) Quantitative analysis of the wound-healing assay as the recovery area in pixels (px) at 24 h. (F) Quantitative analysis of the wound healing assay as the recovery area in pixels (px) after 48 h. All data are presented as mean ± SEM (n = 4 per group); *p < 0.05. (G) Fluorescence microscopy images after EdU proliferation assay showing proliferating cells (Edu-Alexa555; red) and nuclei (DAPI; blue) in NIH/3T3-*Kif3a*-KO cells treated with NIH/3T3-WT-derived and NIH/3T3-*Kif3a*-KO-derived Sup-100K after 48 h of wound development. Scale bar, 125 px. (H) Quantitative analysis of EdU proliferation assay from G. (I) Quantitative analysis of EdU proliferation assay from G. All data are presented as mean ± SEM (N = 3), \**p* < 0.05.

Direct measurement of migration distance further supported these observations. At 12, 24, and 48 h post-treatment, the average migration distance in WT-Sup-100K-treated cells was comparable to that in WT-CM-10K-treated cells, with no statistically significant differences (Figure 2D–F). In contrast, WT-CM-10K induced a ∼1.5-fold increase in migration distance at 12 and 24 h (Figure 2D–E), and a ∼1.9-fold increase at 48 h compared to WT-EV-P3 (Figure 2F), further reinforcing the dominant role of soluble factors over EVs in promoting cell migration.

To assess whether the observed increase in wound healing was also associated with increased proliferation, we performed EdU incorporation assays. *Kif3a*-KO fibroblasts treated with WT-Sup-100K showed no significant difference in EdU labeling compared to those treated with *Kif3a*-KO-Sup-100K in either the wound or non-wound regions at 48h post treatment (Figure 2G). In the wound area, WT-Sup-100K treatment resulted in ∼47% EdU-positive cells, which was not significantly different from the ∼51% observed in cells treated with Sup-100K derived from NIH/3T3-*Kif3a*-KO cells (Figure 2H). Similarly, in the non-wound area, proliferation remained comparable between groups (∼15% vs. ∼19%, respectively; Figure 2I). Notably, a consistent increase in EdU incorporation was observed in the wound area compared to the non-wound region across both treatments, suggesting a general proliferative response to injury that is independent of primary cilia function. These results suggest that the pro-repair effects of WT-Sup-100K are not driven by increased proliferation, but rather by enhanced migratory activity.

These data collectively indicate that soluble bioactive factors present in the 100K × g supernatant of CM from ciliated NIH/3T3 cells are the principal mediators of wound healing in fibroblasts. While EVs might contribute transiently during early stages, the soluble component could predominantly mediate sustained promotion of cell migration.

### Lysophosphatidylcholine (14:0) is a major humoral bioactive factor that increases wound healing in primary cilium-deficient target cells

To identify specific soluble metabolites within the WT-Sup-100K fraction responsible for promoting wound healing in primary cilium-deficient cells, we conducted comparative untargeted metabolomic profiling of Sup-100K fractions derived from NIH/3T3-WT, NIH/3T3-*Kif3a*-KO, and NIH/3T3-*Dync2h1*-KO cell cultures (Figure 3A). A total of 39 metabolites were detected across the samples. Phosphatidylcholines (PCs) comprised the largest class (14 metabolites), followed by sphingomyelins (SMs; 9), lysophosphatidylcholines (LPCs; 8), sphingolipids (SPHs; 4), monoacylglycerols (MGs; 2), and single representatives of methylphosphatidylcholine (MePC) and ceramide (Cer) (Figure 3B; Supplementary Figs. 1-3).

**Figure 3:**
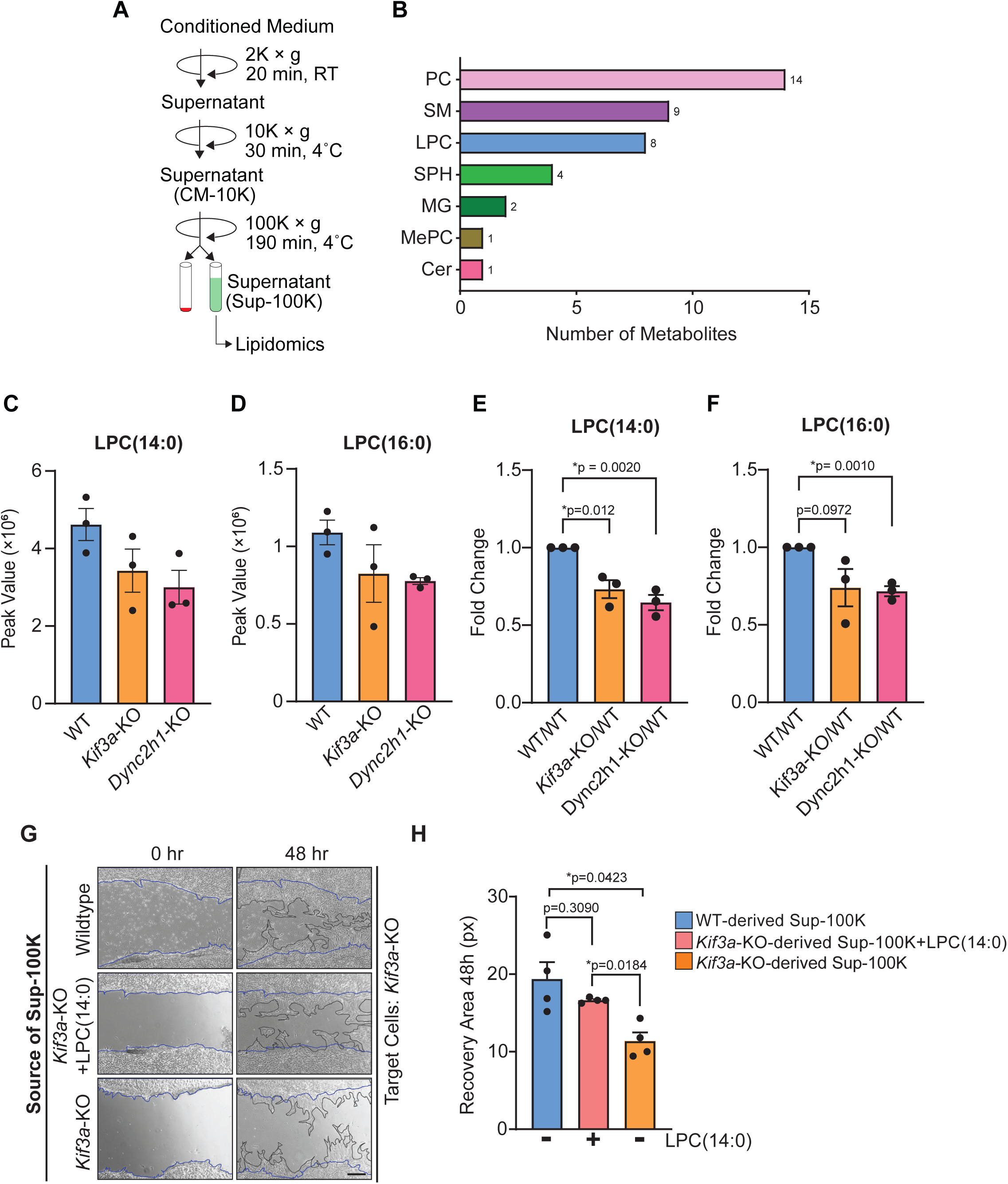
Identification of LPC(14:0) as a major wound-healing bioactive factor in Sup-100K. (A) The diagram demonstrates the centrifugation steps to fraction the 100k-supernatant (Sup-100k) fraction from the conditioned medium for untargeted metabolomics. (B) Number of metabolites and metabolite classes detected in the Sup-100K fractions from NIH/3T3-Wildtype, NIH/3T3-*Kif3a*-KO and NIH/3T3-*Dync2h1*-KO derived conditioned mediums. (C, D) Graphs showing that LPC(14:0) and LPC(16:0) were decreased in both NIH/3T3-*Kif3a*-KO and NIH/3T3-*Dync2h1*-KO vs control depicted as peak values. (E, F) Fold-change decrease in LPC(14:0) and LPC(16:0). (G) Confirmation of the effect of LPC(14:0) on promoting wound healing. Microscopy images of wound closure of *Kif3a*-KO cells treated with *Kif3a*-KO-Sup-100K supplemented with LPC(14:0) (10 μM) (middle panel), WT-Sup-100K (upper panel), and *Kif3a*-KO-Sup-100k (lower panel) at 0 and 48 h after making wound. Blue lines define the area covered by the cells at 0 h after treatment. Dark-grey lines define the area covered by the cells after 48 h. Scale bars, 250 μm. (H) Quantitative analysis of the wound healing assay as the recovery area in pixels (px) at 48 h from G. All data are presented as mean ± SEM (n = 4 per group); \**p* < 0.05.

To pinpoint candidates of bioactive factors that promote wound healing, we focused on metabolites consistently decreased in the Sup-100K fractions from both *Kif3a*-KO and *Dync2h1*-KO cells and increased in the WT-Sup-100K fraction. Among these, two LPC species—LPC(14:0) and LPC(16:0)—were decreased in the cilium-deficient cell lines (Figure 3C-D). Of the two, LPC(14:0) exhibited a significant reduction in both mutant-derived fractions compared to the WT control (Figure 3E-F).

To directly assess the functional role of LPC(14:0) in promoting wound healing, we performed rescue experiments by supplementing *Kif3a*-KO-Sup-100K with 10 µM LPC(14:0). Wound healing responses in primary cilium-deficient target cells were then evaluated and compared with responses to unsupplenmented *Kif3a*-KO-Sup-100K and WT-Sup-100K. Notably, supplementation of LPC(14:0) restored wound closure activity in the *Kif3a*-KO-Sup-100K-treated cells, yielding an average recovery area of ∼17 px, comparable to that induced by WT-Sup-100K (∼20 px), with no statistically significant difference between the two (Figure 3G-H). These data identify LPC(14:0) as a principal humoral bioactive metabolite enriched in the WT-Sup-100K fraction and capable of restoring wound healing capacity in primary cilium-deficient fibroblasts.

### Expression of extracellular matrix-related proteins and cell cycle pathways are regulated by treatment with WT-Sup-100K in primary cilium-deficient target cells

To provide insight on the molecular mechanisms by which WT-Sup-100K promotes sustained wound repair in primary cilium-deficient fibroblasts, we performed transcriptomic profiling of NIH/3T3-*Kif3a*-KO cells treated with either WT-Sup-100K or *Kif3a*-KO-Sup-100K for 48 hours. RNA sequencing followed by differential gene expression analysis (adjusted *p* < 0.05) revealed 439 significantly altered genes, the majority (n = 408) of which exhibited modest changes in expression (|log_2_ fold change| < 0.5; *q* ≤ 0.05) (Figure 4A, green). A subset of 31 genes met both statistical and biological significance criteria (|log_2_ fold change| > 0.5; adjusted *p* < 0.05), of which 6 were upregulated (Figure 4A, red) and 25 downregulated (Figure 4A, blue) in response to WT-Sup-100K treatment relative to control.

**Figure 4:**
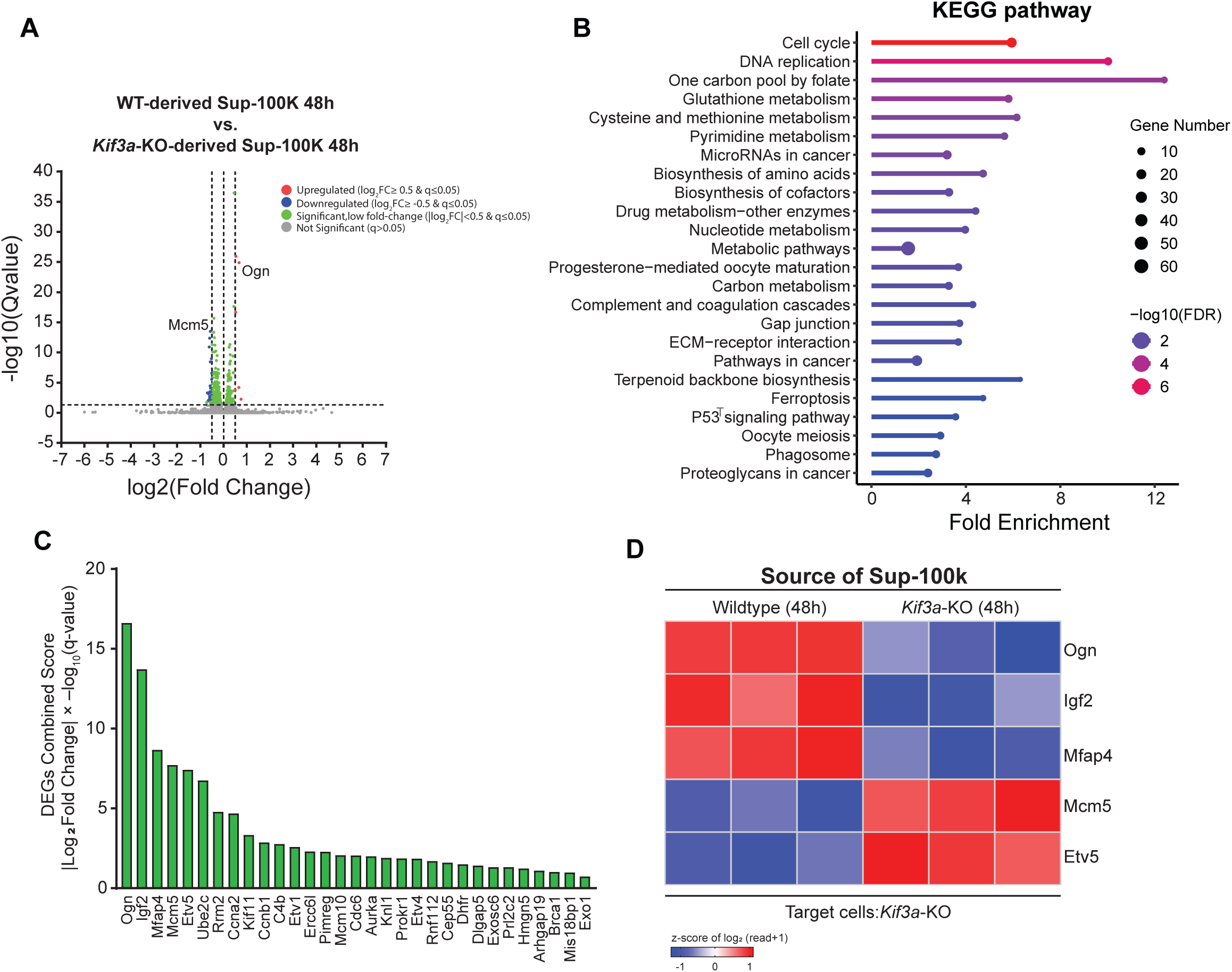
Transcriptomic profiling and functional enrichment analysis of differentially expressed genes (DEGs) in target cells treated with Sup-100K fraction. (A) Volcano plot displaying 439 DEGs between WT-derived Sup-100K–treated and *Kif3a*-KO-derived Sup-100K–treated cilium-deficient fibroblasts. The x-axis shows log_2_ fold change, and the y-axis shows –log_10_(Q-value). Vertical dashed lines represent the fold change threshold (|log_2_FC| ≥ 0.5), and the horizontal dashed line denotes the significance cutoff (–log_10_(q) ≥ 1.3). Red and blue points mark DEGs meeting both thresholds, green points indicate genes significant only by q-value, and grey points are non-significant. (B) KEGG pathway enrichment analysis of all 439 DEGs, with significantly enriched pathways ranked by combined score (–log_10_(FDR) × fold enrichment). (C) Bar graph of combined scores for 31 DEGs selected based on biological significance, integrating fold change magnitude and statistical significance (– log_10_(q-value)). Higher scores reflect stronger combined significance. (D) Heat map of the top 5 DEGs from panel C, showing z-score–normalized expression levels across individual replicates for WT-derived Sup-100K–treated and *Kif3a*-KO-derived Sup-100K–treated cells. Red indicates higher expression, and blue indicates lower expression relative to the mean.

To identify the broader biological pathways impacted by WT-Sup-100K, we performed KEGG pathway enrichment analysis using all significant DEGs (q ≤ 0.05), independent of fold-change magnitude, in order to capture transcriptional shifts with potential cumulative effects. This analysis revealed significant enrichment in diverse pathways, including those associated with extracellular matrix dynamics (*ECM-receptor interaction*, *Proteoglycans in cancer*), cell cycle regulation (e.g., *Cell cycle*, *DNA replication*, *p53 signaling pathway*), and metabolic processes (e.g., *One carbon pool by folate*, *Carbon metabolism*, *Glutathione metabolism*, *Pyrimidine metabolism*) (Figure 4B). Additional enriched categories included *Gap junction*, *Ferroptosis*, *Phagosome*, *MicroRNAs in cancer*, and *Biosynthesis of cofactors*, highlighting the broad cellular remodeling induced by WT-Sup-100K.

To prioritize the most biologically relevant targets, we computed a combined score for each DEG by multiplying the absolute log_2_ fold change by the –log_10_(q-value), integrating both effect size and statistical strength. The five highest-ranking genes by this metric were *Osteoglycin* (*Ogn*), *Insulin-like growth factor 2* (*Igf2*), *Microfibril-associated glycoprotein 4* (*Mfap4*), *Minichromosome maintenance complex component 5* (*Mcm5*), and *Ets variant 5* (*Etv5*), with combined scores of 16.63, 13.73, 8.67, 7.71, and 7.42, respectively (Figure 4C). These genes were then visualized in a heatmap using row-wise z-score normalization across biological replicates (Figure 4D).

Among the top five, *Ogn*, *Igf2*, and *Mfap4* were upregulated in response to WT-Sup-100K (Figure 4D). Notably, *Ogn* and *Mfap4* encode extracellular matrix (ECM)-associated proteins, consistent with enhanced migratory behavior observed in functional assays. *Ogn* encodes a small leucine-rich proteoglycan implicated in ECM remodeling, while *Mfap4* contributes to matrix organization and fibroblast motility. In contrast, *Mcm5* and *Etv5*, both key regulators of cell cycle and DNA replication, were downregulated following WT-Sup-100K treatment (Figure 4D). *Mcm5* plays an essential role in DNA replication licensing, while *Etv5* functions as a transcription factor involved in various development and cellular processes. This transcriptomic signature suggests that WT-Sup-100K promotes a shift toward a matrix-remodeling, pro-migratory state in cilium-deficient fibroblasts.

Given the modest yet measurable early-stage wound healing effects of the WT-EV-P3 fraction, we also examined the proteomic response in *Kif3a*-KO cells following a short-term treatment with a highly concentrated EVs fraction isolated via ultracentrifugation (UC) or ultrafiltration (UF) (Supplementary Figures 4A and 5A). Twelve hours post-treatment, both EV preparations, WT-EV-UC and WT-EV-UF, upregulated expression of the basement membrane glycoprotein *Nidogen-2* (Nid2), a key structural component of the ECM (Supplementary Figures. 4B and 4D and 5B and 5D). This suggests that EV-associated signals could contribute to early ECM remodeling events that support migration during the initial phase of wound repair.

Together, these transcriptomic analyses imply that WT-Sup-100K promotes wound healing in primary cilium-deficient fibroblasts through coordinated activation of ECM-associated protein expression and cell cycle signaling. Additionally, EV-mediated regulation of ECM modulators such as Nid2 could further facilitate the early stages of wound healing.

## Discussion

This study uncovers a previously underappreciated function of the primary cilium in coordinating wound repair via regulating release of humoral bioactive factors that act on target cells through paracrine signaling. While the primary cilium has classically been defined as cellular antenna coordinating intracellular responses to extracellular cues, our data demonstrate that it also governs the secretion of bioactive factors capable of influencing neighboring/distant cells. Specifically, we show that ciliated fibroblasts secrete soluble factors that promote migration in primary cilium-deficient fibroblasts, revealing a paracrine-like mechanism that depends on the donor cell’s ciliary status. This finding extends emerging evidence that the primary cilium can regulate not only intracellular signaling but also paracrine and endocrine outputs. A previous work by Hu et al. demonstrated a cilium-dependent mode of ciliary disassembly through LPA secretion, while Verbruggen et al. recently identified a cilium-regulated paracrine axis in osteocytes that suppresses tumor cell proliferation via TNF-α^[29,30]^. Our study adds to this growing body of evidence by showing that ciliated fibroblasts secrete humoral factors capable of enhancing wound repair in a non-cell-autonomous fashion.

Our observation that WT-CM-10K enhanced wound recovery in two independent cilium-deficient lines, *Kif3a*-KO (which lacks anterograde intraflagellar transport) and *Dync2h1*-KO (which lacks retrograde transport) (Figure 1), argues against clone-specific effects and instead supports a general requirement for intact ciliogenesis in promoting a wound healing secretory profile. The fact that WT-CM significantly improved both the area and the distance of recovery in the target cells further supports the notion that primary cilia influence the release of functional humoral factors, rather than simply altering baseline fibroblast motility.

Another important observation in this study is that conditioned medium (CM) from primary ciliated fibroblasts, but not from primary cilium-deficient counterparts promotes wound closure in *Kif3a*-KO fibroblasts. This suggests that the primary cilium as a regulator of secretome composition, particularly in the context of wound repair. We show that this effect is attributable to the soluble fraction of the CM, as removal of EVs by ultracentrifugation does not diminish the wound healing activity. This observation suggests that small bioactive molecules rather than EVs are the principal mediators of the effect.

Notably, despite the increase in migratory activity in WT-Sup-100K-treated cilium-deficient fibroblasts, proliferation rates remained unchanged compared to cells treated with *Kif3a*-KO-Sup-100K. EdU incorporation assays revealed that proliferation within the wound area was approximately three times higher than in non-wound regions for both WT- and *Kif3a*-KO-derived 100k supernatant-treated cells, indicating a robust proliferative response localized to the wound site. However, the lack of significant difference in proliferation between these treatment groups suggests that the enhanced wound healing observed is primarily driven by increased migration rather than cell division. This points to a primary cilium-independent mechanism for the proliferative response during wound repair, while cilium-dependent secreted factors specifically promote motility to accelerate closure.

By applying untargeted metabolomic profiling, we identified LPC(14:0) as a key component enriched in CM from primary ciliated cells but depleted in CM from primary cilium-deficient cultures. Supplementing *Kif3a*-KO-CM with LPC(14:0) restored wound-healing activity in target cells, supporting a causative role for this metabolite (Figure 3). LPCs are bioactive lipids with a variety of functions, typically produced from cell membranes as normal metabolic products of phosphatidylcholine (PC) ^[52]^. Furthermore, these lipids can be released into the extracellular environment as bioactive factors. For example, stimulated bronchial epithelial cells release bioactive LPCs 16:0, 18:0, and 18:1 in response to VEGF^[53]^. The ability of LPCs to restore wound healing in *Kif3a*-KO target cells aligns with previous studies, highlighting their role in enhancing cell migration. For instance, ID8 cells stimulated with LPC showed increased cell migration and invasion ^[54]^. Similarly, LPC also induced the migration of human coronary artery smooth muscle cells through the release of endogenous bFGF.^[55]^

The enrichment of LPC(14:0) in CM from primary ciliated cells raises the possibility that the primary cilium modulates not only signaling pathways but also lipid biosynthesis and secretion. Indeed, the ciliary membrane possesses a distinct lipid composition^[10,56–59]^ and hosts several lipid-modifying enzymes^[60,61]^, including phosphoinositide phosphatases and cholesterol transporters^[62,63]^. It is conceivable that disruption of ciliary structure or function leads to aberrant lipid processing, thereby altering the extracellular signaling landscape. This idea is supported by metabolomic studies in human ciliopathies such as Bardet-Biedl and Alström syndromes, which are characterized by systemic metabolic disturbances, including reduced serum levels of LPCs^[64]^. Our data suggest a possible mechanistic basis for these observations, wherein loss of ciliary control over lipid metabolism impairs the cell’s ability to produce and secrete key signaling molecules such as LPC(14:0).

Beyond identifying the active metabolite, our transcriptomic analysis of primary cilium-deficient target cells exposed to WT-Sup-100K provides further insight into downstream mechanisms. We observed the activation of a gene expression program enriched for ECM-related proteoglycans. Notably, expression of genes such as *Ogn*, *Mfap4*, and *Igf2* was upregulated, all of which have documented roles in promoting tissue remodeling and regeneration. For instance, *Ogn*, a small leucine-rich proteoglycan, is known to promote epithelial repair and support neurite outgrowth via IGF-2 signaling^[65–67]^. *Mfap4*, a component of elastic fiber microfibrils, facilitates cell adhesion and has been associated with migratory behavior in lung adenocarcinoma cells^[68]^. Although classically regarded as a mitogen, *Igf2* also promotes fibroblast and epithelial motility through activation of PI3K-Akt and MAPK signaling pathways, which are essential for cytoskeletal dynamics and directional migration^[69–71]^. Conversely, we observed downregulation of *Mcm5* and *Etv5* expression in WT-Sup-100K–treated fibroblasts. While *Mcm5* is canonically involved in DNA replication, its overexpression has been shown to impair endodermal migration in the zebrafish model through a Cxcr4a–Itgb1b–dependent mechanism, independent of cell cycle regulation^[72]^. Etv5, a transcription factor typically associated with pro-migratory roles in EMT and cancer progression^[73]^, was paradoxically suppressed in our system despite increased cell motility. This implies that Etv5 might play a context-dependent role, and its downregulation could reflect a distinct migratory program in mesenchymal fibroblasts undergoing collective wound closure. The upregulation of ECM-associated genes expression suggests that WT-Sup-100K reprograms the wound microenvironment in target cells to support both structural remodeling and cellular motility.

Interestingly, although EVs were not the dominant mediators of sustained wound healing, proteomic analysis indicated that they could contribute to early remodeling events. Specifically, a highly concentrated EVs fraction from ciliated donor cells induced expression of Nidogen-2, a key ECM glycoprotein involved in basement membrane integrity^[74,75]^. This suggests a model in which EVs deliver structural cues that facilitate initial migration, while soluble metabolites such as LPC sustain longer-term proliferation and wound closure. Temporal coordination between vesicle-bound and soluble molecules could therefore underlie the full wound healing capacity of the fibroblast secretome.

This work has several broader implications. First, it reveals that ciliated cells can exert influence over neighboring/distant cells through noncanonical secretory pathways that extend beyond morphogen signaling. This has potential relevance for tissue homeostasis and regeneration in ciliated tissues such as the kidney, lung, and skin, where fibroblasts and epithelial cells must coordinate responses to injury. Second, the apparent involvement of ECM-related gene expression and remodeling suggests that the primary cilium could indirectly regulate the composition and mechanical properties of the tissue microenvironment.

Nevertheless, several questions remain. The upstream pathways by which the primary cilium regulates LPC production are unclear and would involve both transcriptional and post-translational mechanisms. Additionally, while LPC(14:0) was sufficient to restore wound healing capacity, it is unlikely to be the sole effector. Other low-abundance lipids, peptides, or nucleotides might contribute synergistically, but failed to be detected due to technical limitations or serum interference. Finally, it will be important to determine whether this secretory axis is conserved *in vivo* and in other cell types in the future work.

In conclusion, this study identifies a primary cilium-dependent paracrine signaling mechanism in fibroblasts that promotes wound repair through soluble factors. It reveals that the primary cilium plays a central role not only in sensing extracellular signals but also in shaping the extracellular environment to direct wound repair. These findings open new avenues for investigating primary cilium-mediated intercellular communication and may have translational relevance for regenerative medicine and ciliopathy-related disorders.

## Acknowledgments

This work was supported in part by the Japan Society for Promotion of Science Grants-in-Aid for Scientific Research Activity Start-up 19K23728 and for Early-Career Scientists 21K15088 (to Faryal Ijaz), and the Japan Science and Technology Agency, Precursory Research for Embryonic Science and Technology JPMJPR17H1 (to Koji Ikegami) and JPMJPR18H2 (to Koshi Imami). Part of this work was conducted at the Natural Science Center for Basic Research and Development, Hiroshima University. We thank Ms. Madoka Hamada and Dr. Qushay Umar Malinta for technical assistance.

## Author contributions

Koji Ikegami and Faryal Ijaz designed research; Faryal Ijaz, and Koshi Imami performed experiments and acquired the data; Faryal Ijaz, Koshi Imami, and Koji Ikegami analyzed the data; Faryal Ijaz and Koji Ikegami wrote the paper; all authors discussed the results and commented on the manuscript.

## Additional Information

Competing Interests: The authors have no competing interests to declare.

## Data Availability

Data generated in this study is available upon request.

---

**Supplementary Figure 1: Remaining results of metabolomics analyses of Sup-100K.** (A-N) Graphs showing comparisons of different PCs detected in NIH/3T3-*Kif3a*-KO and NIH/3T3-*Dync2h1*-KO vs control are depicted as peak values. (O) Graph showing comparison of MePC(35:0) detected in NIH/3T3-*Kif3a*-KO and NIH/3T3-*Dync2h1*-KO VS control depicted as peak values.

**Supplementary Figure 2: Remaining results of metabolomics analyses of Sup-100K.** (A-I) Graphs showing comparisons of different SMs detected in NIH/3T3-*Kif3a*-KO and NIH/3T3-*Dync2h1*-KO vs control are depicted as peak values. (J-M) Graphs showing comparisons of different SPHs detected in NIH/3T3-*Kif3a*-KO and NIH/3T3-*Dync2h1*-KO vs control are depicted as peak values. Graphs comparing MG(18:0) and MG(18:0)(rep) detected in NIH/3T3-*Kif3a*-KO and NIH/3T3-*Dync2h1*-KO vs control are depicted as peak values. (O) Graph showing comparison of Cer (d34:0) detected in NIH/3T3-*Kif3a*-KO and NIH/3T3-*Dync2h1*-KO vs control depicted as peak values.

**Supplementary Figure 3: Remaining results of metabolomics analyses of Sup-100K.** (A-H) Graphs showing comparison of different LPCs detected in NIH/3T3-*Kif3a*-KO and NIH/3T3-*Dync2h1*-KO vs control are depicted as peak values.

**Supplementary Figure 4: Proteomic analysis of target cells treated with EV-UC fraction. (**A) Schematic representation of the fractionation of the EV fraction (EV-UC) from conditioned medium after ultracentrifugation. (B) Scatter plot analysis of the proteome profiles of WT-EV-UC12h vs *Kif3a*-KO-EV-UC 12h treatment in target cells. Red and green dots highlight significantly upregulated and downregulated genes, respectively. (C) Statistics of enriched GO terms are shown in Panel B. The size of the point indicates the number of DEGs in this pathway, and the color of the points corresponds to the *p*-value range. (D) Heat maps showing proteins detected in each sample represented as z-transformed log2 (LFQ intensity) from A.

**Supplementary Figure 5: Proteomic analysis of target cells treated with the EV-UF fraction. (**A) Schematic representation of the fractionation of the EV fraction (EV-FC) from conditioned medium after 100 K ultrafiltration. (B) Scatter plot analysis of the proteome profiles of WT-EV-UF 12h vs *Kif3a*-KO-EV-UF 12h treatment in target cells. Red and green dots highlight significantly upregulated and downregulated genes, respectively. (C) Statistics of enriched GO terms are shown in Panel B. The size of the point indicates the number of DEGs in this pathway, and the color of the points corresponds to the *p*-value range. (D) Heat maps showing proteins detected in each sample represented as z-transformed log2 (LFQ intensity) from A.

