## Supplementary figures and images for "Primary Cilium-dependent Humoral Bioactive Factors Acts in a Paracrine Manner to Control Fibroblast Cell Migration"

### Supplementary Figure 1

#### Supplementary Figure 1

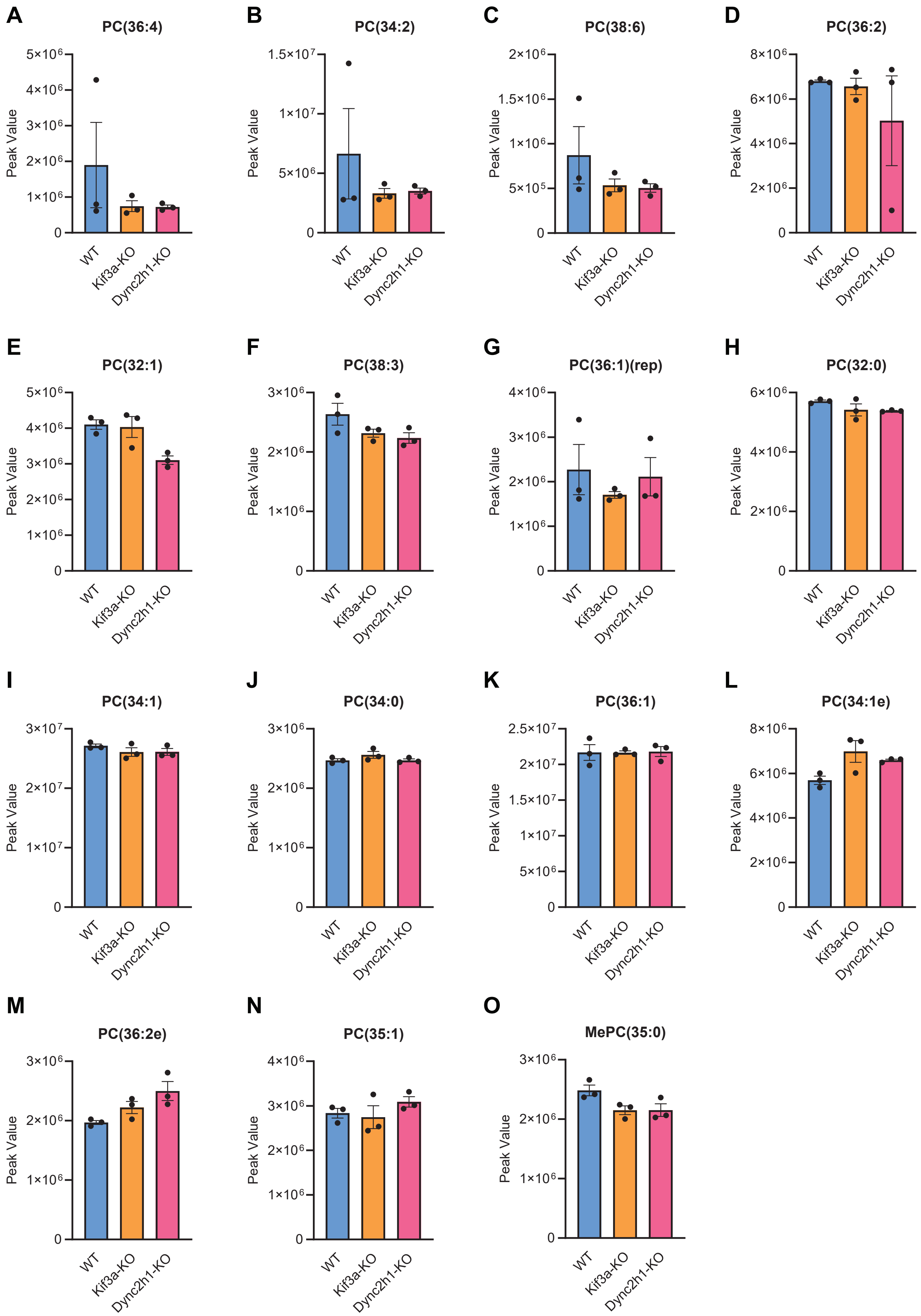

### Supplementary Figure 2

#### Supplementary Figure 2

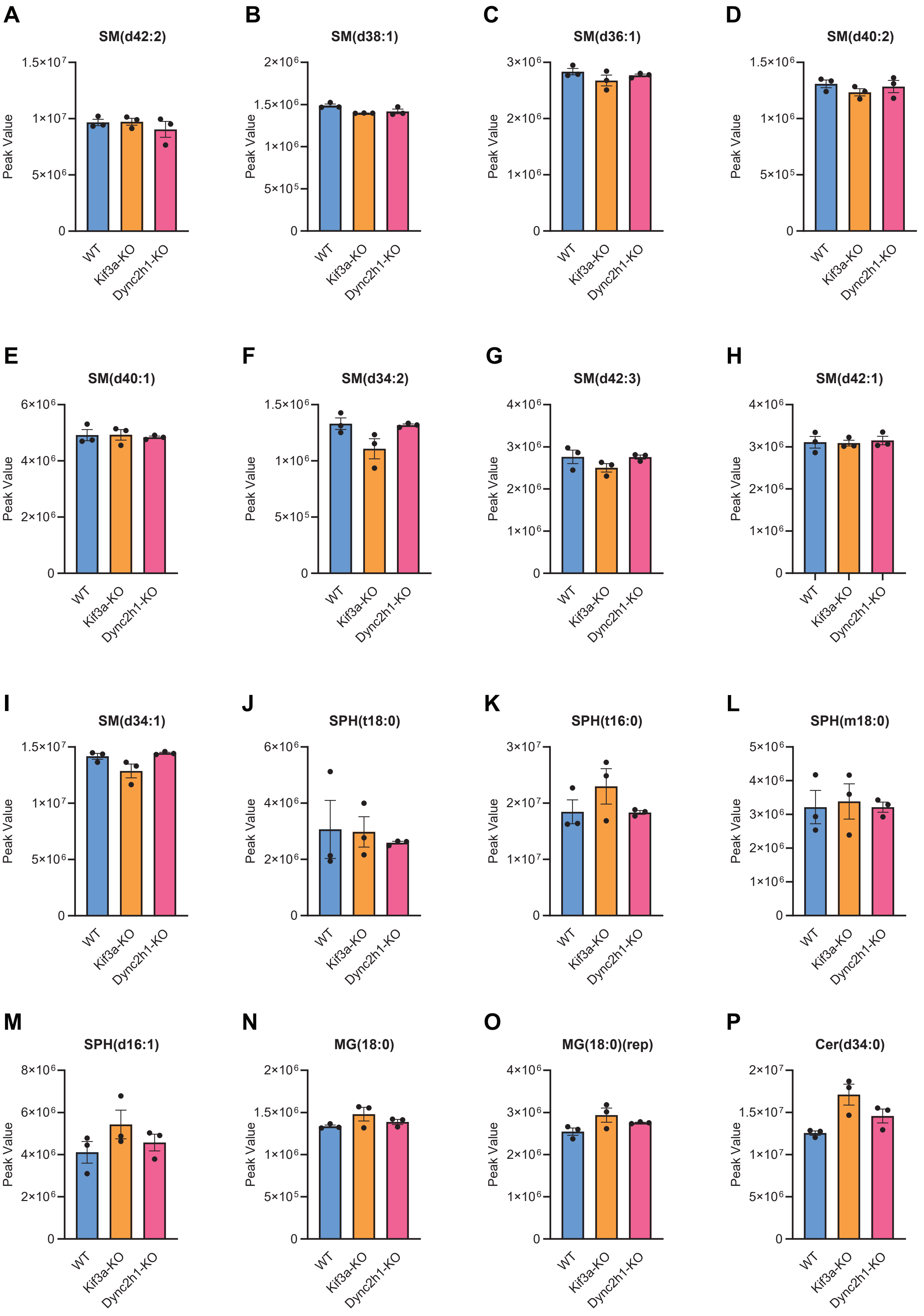

### Supplementary Figure 3

#### Supplementary Figure 3

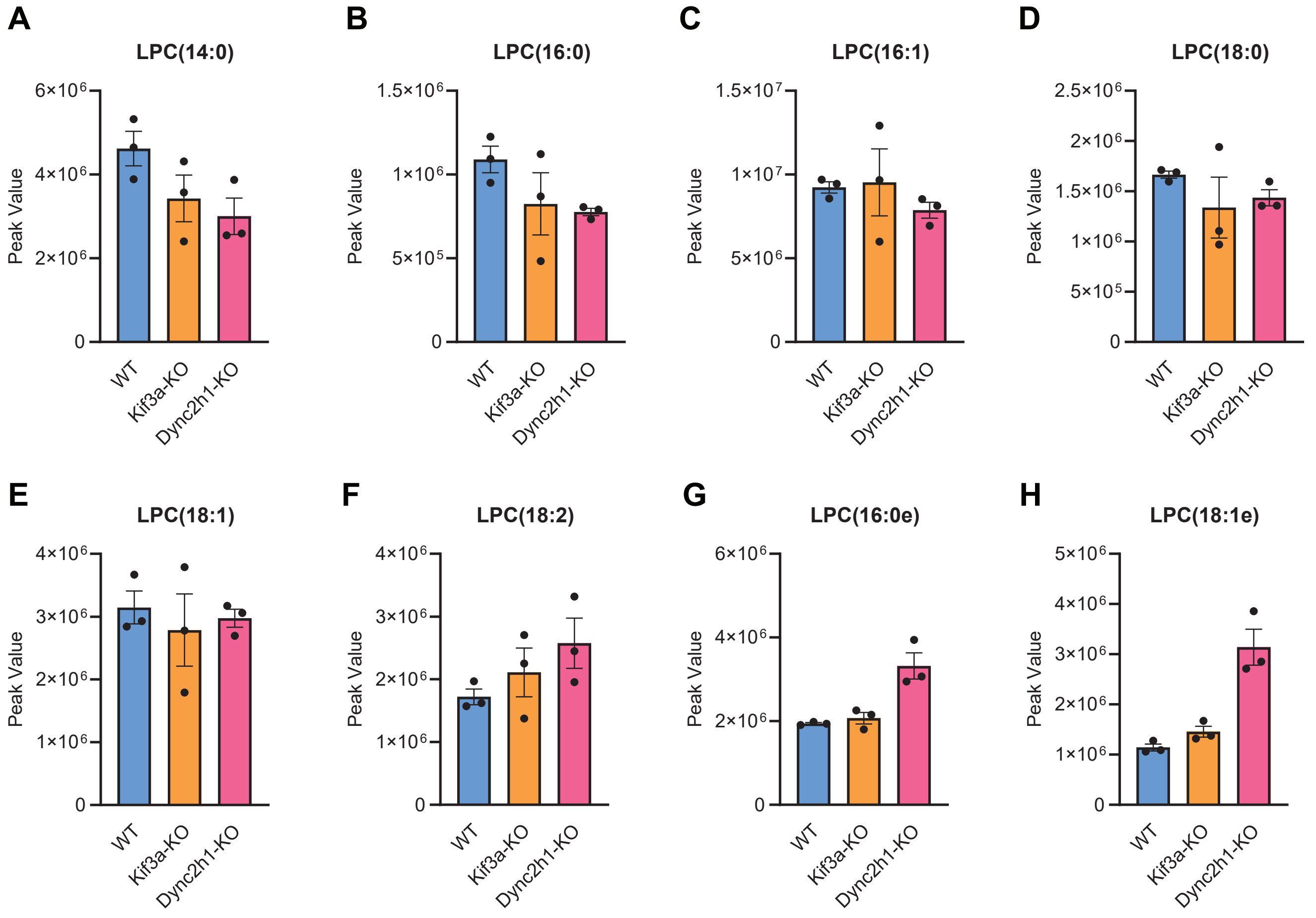

### Supplementary Figure 4

#### Supplementary Figure 4

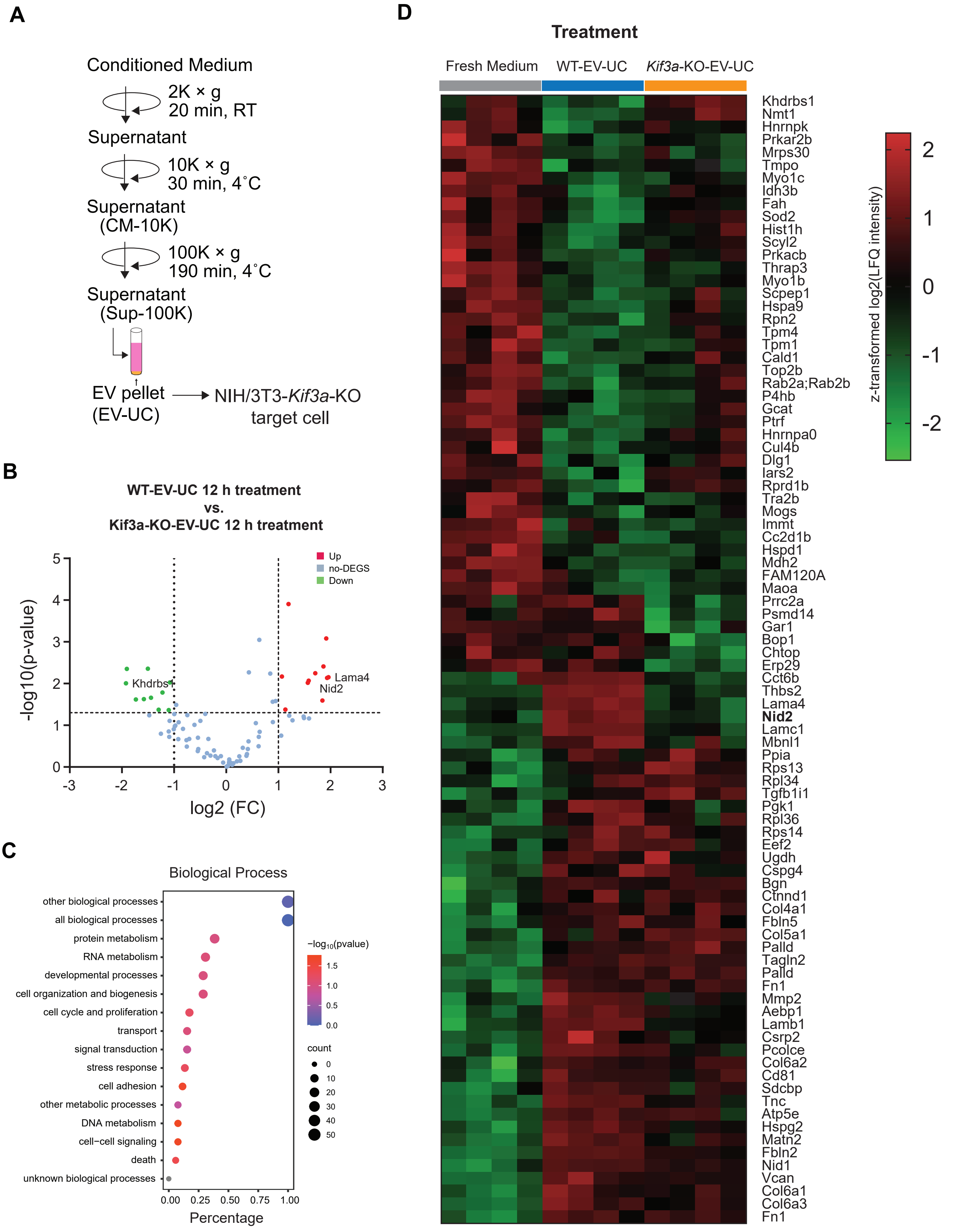

### Supplementary Figure 5

#### Supplementary Figure 5

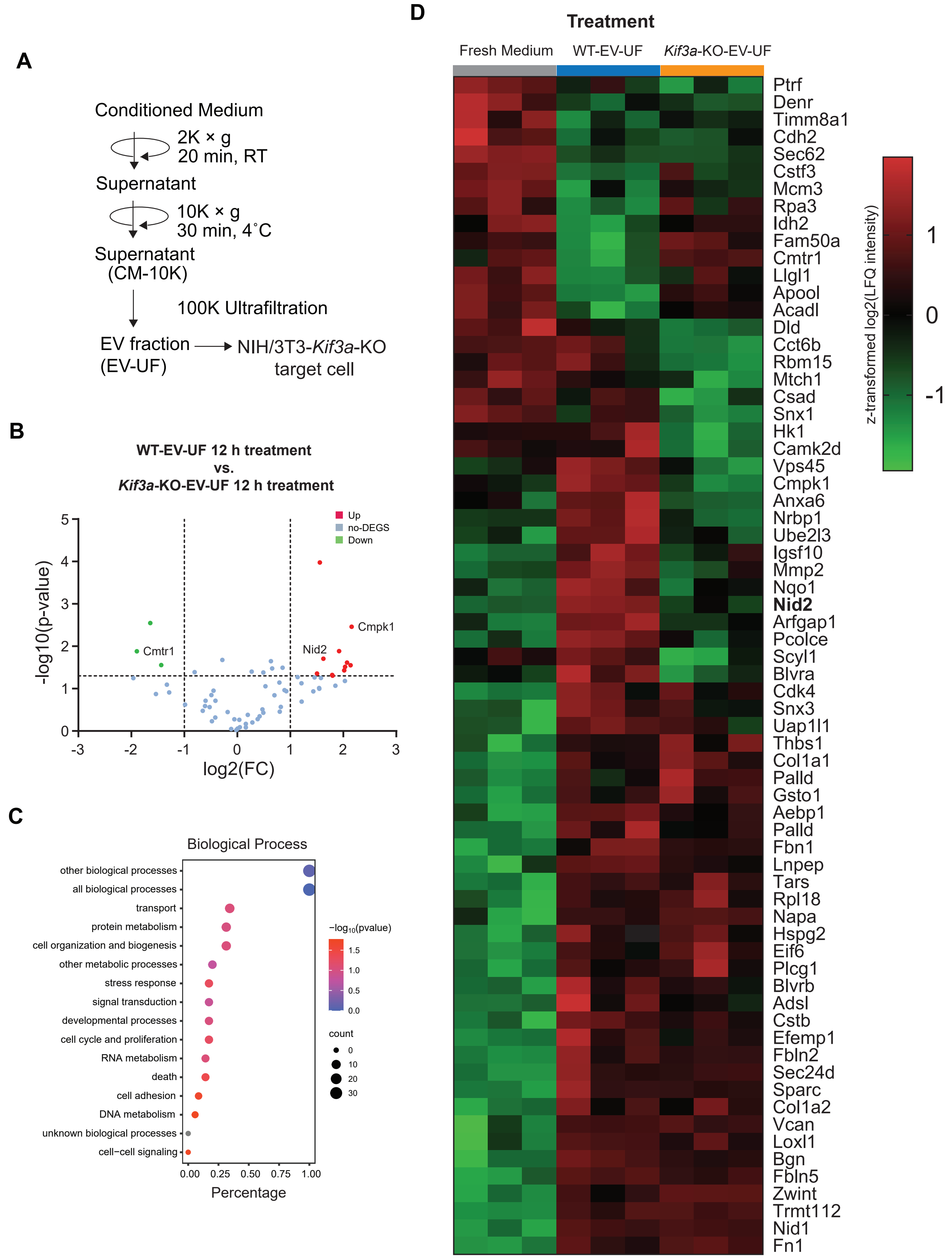
